# Interferon signaling promotes early neutrophil recruitment after zebrafish heart injury

**DOI:** 10.1101/2025.11.16.688717

**Authors:** Alexis V. Schmid, James A. Gagnon

## Abstract

Heart injury triggers a robust cellular response in zebrafish, characterized by neovascularization, immune cell activation, and the infiltration of proliferative cardiomyocytes that collectively lead to scarless regeneration. Upon injury, damage-associated molecular patterns are released by dying cells and injured extracellular matrix. These molecules bind to pattern recognition receptors on various cell types, promoting inflammation and immune cell recruitment by upregulating chemokines and pro-inflammatory cytokines. We previously identified the activation of injury-induced interferon signaling as a distinguishing feature between the regenerative zebrafish and non-regenerative medaka. Here, we establish interferon-Φ1 (IFN□1) as the primary driver of interferon signaling after zebrafish heart injury. IFN□1 expression is induced hours after injury and directs interferon-stimulated gene expression, which peaks at 72 hours post-injury. This response is lost in *ifnphi1* mutants, which disrupt IFN□1 expression. *ifnphi1* mutants have reduced neutrophil recruitment to the injured myocardium, while macrophages and neovascularization are unaffected. By studying later stages of regeneration, we find that *ifnphi1* mutant hearts have a modest defect in scar resolution. Collectively, these findings uncover a critical early signaling cascade during the inflammatory response to heart injury and provide new insights into the mechanisms that choreograph zebrafish heart regeneration.

## INTRODUCTION

Myocardial infarction occurs when blood flow to a portion of the heart is blocked, leading to irreparable cell death and the formation of a large, inelastic scar that hinders heart function (Salari et al., 2023). Nearly all mammals lose their regenerative capacity after the neonatal stage, whereas most fish and amphibians retain it throughout development and into adulthood. These regenerators replace collagenous scar tissue with fresh cardiomyocytes to restore heart function to the injured myocardium (González-Rosa et al., 2011; González Rosa et al., 2017; Poss et al., 2002). Zebrafish have emerged as a powerful model to study cardiac regeneration (Gemberling et al., 2013). Following cardiac injury, zebrafish employ a sequence of pro-inflammatory and anti-inflammatory immune signals to coordinate collagen deposition, revascularization, and eventual scar resolution. In contrast, the non-regenerating medaka fish exhibit a prolonged pro-inflammatory immune response after injury, leading to scar persistence (Lai et al., 2017). Studies of regenerators may yield new therapeutic insights into the specific immune and cellular mechanisms required to resolve the scar and regenerate functional cardiac tissue.

In zebrafish, heart regeneration is initiated by a choreographed immune response. The initial injury triggers the release of damage-associated molecular patterns (DAMPs) from necrotic tissue and injured extracellular matrix. These DAMPs induce inflammation by binding to pattern recognition receptors (PRRs) on multiple cell types to recruit waves of immune cells to the wound via the release of pro-inflammatory cytokines and chemoattractants (Iribarne, 2021; Julier et al., 2017). Neutrophils are the first pro-inflammatory cells recruited to the injury zone, rapidly localizing to the wound as early as six hours post-injury (hpi) (Julier et al., 2017). Neutrophils are vital for wound detection by helping to recruit monocytes, which differentiate into macrophages at the wound site (Feng et al., 2025; Wilgus et al., 2013). In addition to promoting inflammation early after injury, neutrophils also secrete vascular endothelial growth factor (VEGF-A), which promotes early neovascularization of the injured myocardium (El-Sammak et al., 2022; Gong & Koh, 2010; Lörchner et al., 2023). Neutrophils also activate epicardial cells to induce epithelial-to-mesenchymal transition, enabling epicardial infiltration and significant contributions to wound healing (Peterson et al., 2024).

Macrophages are the second immune cells recruited to the injury site, providing both a pro-inflammatory and an anti-inflammatory response (Bevan et al., 2020; Sánchez-Iranzo et al., 2018). The numerous roles of macrophages are still being explored; however, we know they shape the inflammatory response by phagocytosing pro-inflammatory apoptotic neutrophils, clearing cellular debris, shaping new extracellular matrix and neovasculature, and depositing collagen alongside fibroblasts for scar formation (Bevan et al., 2020; Constanty et al., 2025; Goumenaki et al., 2024; Julier et al., 2017). While immune cells are required for complete regeneration, the specific immune responses that distinguish regenerating and non-regenerating species are still being investigated (Lai et al., 2017; Peterson et al., 2024).

We previously compared the response to heart injury in two distantly related teleost species, medaka and zebrafish. Despite these fish having similar heart morphology and gene orthology, medaka cannot recover injured myocardium. The innate immune landscape in medaka is different from that of zebrafish, with higher neutrophil recruitment, less macrophage recruitment, and an overall more pro-inflammatory response (Lai et al., 2017). Interestingly, stimulating macrophage recruitment with the TLR agonist, Poly I:C, stimulates heart regeneration in medaka, indicating a failure in the initial damage-sensing phase (Lai et al., 2017). However, the specific signaling pathways underlying this process in zebrafish remain unclear. Using single-cell RNA sequencing, we identified a key difference in the inflammatory response after injury. We found that medaka lacked an injury-induced interferon signaling response to injury. Conversely, zebrafish display robust upregulation of interferon-stimulated genes (ISGs) across most cardiac cell types at three days post-injury (dpi) during the inflammatory stage (Carey et al., 2024).

Interferons are pro-inflammatory cytokines that are released by immune cells responding to infection or injury (Cheng et al., 2008). There are three types of interferons in zebrafish: Type I (IFNΦ1-4), Type II (IFNγ1-2), and Type IV (IFNυ) (Aggad et al., 2009; S. N. Chen et al., 2022; Cheng et al., 2008). IFNγ studies in mice and in cell culture have highlighted the importance of Type II interferon signaling in macrophage activation and successful skeletal muscle regeneration. However, prolonged IFNγ exposure in these models led to excessive fibrosis, reduced muscle contractility, and inhibited regeneration (Adhikari et al., 2020; Z. Chen et al., 2021; Cheng et al., 2008). Our single-cell RNA sequencing analysis of injured zebrafish and medaka hearts revealed specific upregulation of *ifnphi1* in injured zebrafish hearts (Carey et al., 2024). Still, the role of Type I interferon signaling in heart regeneration remains unexamined. Interferons bind to various cell types in the heart, which leads to the expression of ISGs. ISGs have defined roles in pathogen clearance by recruiting innate immune cells to the infection (Balla et al., 2020; Schneider et al., 2014); however, whether ISGs play a role in coordinating a pro-regenerative immune response during heart regeneration is unclear.

Here, we establish IFN 1 as the principal driver of interferon signaling following heart injury in zebrafish. *ifnphi1* gene expression is rapidly induced at the injury site, pointing to a role in early damage sensing and immune cell recruitment. We find that IFN 1 is required to recruit neutrophils to the injury site, an important early step for creating a pro-regenerative environment. Despite this key role in early inflammation, *ifnphi1^-/-^* showed no defects in later regenerative processes, including macrophage recruitment and neovascularization. However, injured *ifnphi1^-/-^* hearts still had a modest defect in scar resolution. Overall, *ifnphi1*-driven interferon signaling orchestrates a discrete, essential role in the initiation of inflammation without being requisite for the subsequent complete regeneration of the zebrafish heart.

## RESULTS

### Interferon signaling is an immediate and injury-specific pathway activated during the inflammatory response

We previously identified interferon-stimulated genes (ISGs) as significantly upregulated 3 dpi in zebrafish endocardial, epicardial, and leukocyte cell populations within the ventricle, and coincide with an increase in *ifnphi1* positive cells (Carey et al., 2024). In contrast, non-regenerative medaka lack an injury-induced interferon response, suggesting that interferon signaling may play an important role in creating a pro-regenerative environment in zebrafish early after heart injury. To better characterize this signaling pathway in zebrafish, we examined transcript levels of *ifnphi1* mRNA in zebrafish heart ventricles in a time course following experimental cryoinjury (Fig.1A). Relative to the uninjured control hearts, *ifnphi1* expression peaked 2 hours after cryoinjury and returned to baseline expression levels by 24 hpi (Fig. 1A). *ifnphi*2-4 transcript levels were unchanged in injured hearts relative to uninjured hearts driving our focus on investigating *ifnphi1* (Fig. 1B). Given the role of interferon signaling in pathogen clearance, and the common presence of infectious agents in fish facility water (Balla et al., 2020), we tested for the upregulation of *ifnphi1* after sham injury. This injury exposes the fish’s chest cavity to the fish facility water without damaging the ventricle. Here, no upregulation of *ifnphi1* was observed, providing evidence that interferon signaling is specifically responding to heart cryoinjury and not chest cavity exposure alone (Fig. 1C).

**Figure 1:**
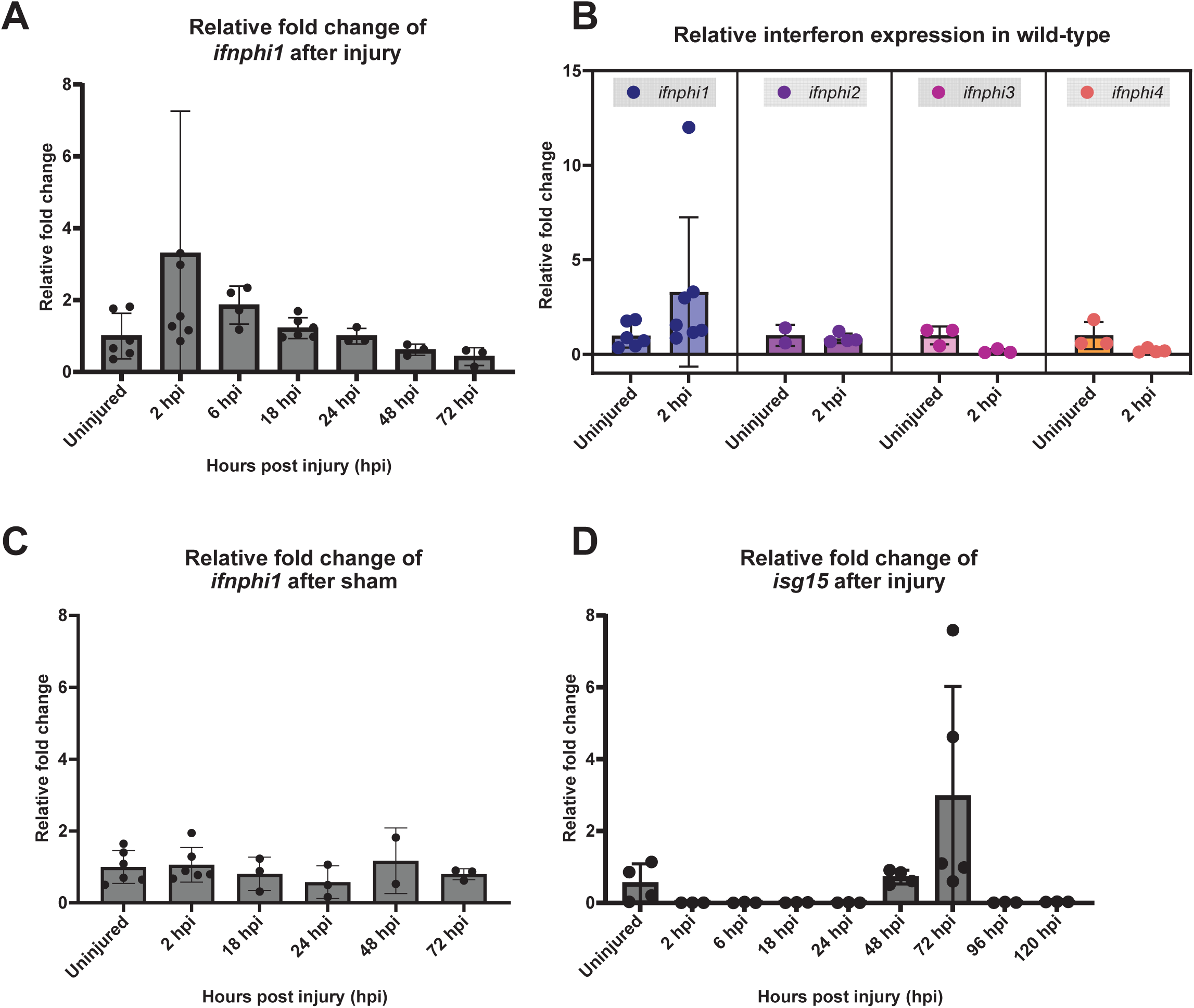
Interferon signaling is induced by zebrafish heart injury. A) A time course of *ifnphi1* gene expression after cryoinjury (2 to 72 hpi) normalized to *elf1a* expression. Relative fold change calculated by normalizing to uninjured control ventricles. B) *ifnphi1-4* gene expression in wildtype ventricles at 2 hpi relative to uninjured ventricles. C) A time course of *ifnphi1* gene expression after sham injury (2 to 72 hpi) relative to uninjured ventricles. D) A time course of *isg15* gene expression after cryoinjury (2 to 120 hpi) relative to uninjured ventricles. Each dot shown represents a single ventricle after dissection away from the atrium and bulbus arteriosus. Data are shown as the mean ±□s.d.

To further characterize the production of ISGs downstream of interferon signaling, we used RT-qPCR to measure the relative fold change of *interferon-stimulated gene-15* (*isg15*) mRNA over an early injury time course in wild-type zebrafish. Relative to uninjured hearts, *isg15* expression peaks at 72 hpi, well after the initial expression of interferon mRNA (Fig. 1D). Together, these data describe the tempo of interferon signaling after zebrafish heart injury: induced immediately after injury and received by endocardial, epicardial, and leukocyte cells to promote the expression of interferon-stimulated genes within the first three days after injury.

### Generating an *ifnphi1* mutant zebrafish

To study the role of interferon signaling in heart regeneration, we generated an *ifnphi1^-/-^* zebrafish. Wildtype embryos were injected with four gRNAs targeting exon 1 and exon 3 of the *ifnphi1* gene. Through crossing we established a stable line with a large deletion between these target sites which is predicted to disrupt protein function (Fig. 2A and Supplementary Table). We assessed the functional disruption of interferon signaling in the mutant using RT-qPCR to evaluate the expression of *isg15* in injured mutant and wild-type heart ventricles. *ifnphi1^-/-^* fish had no induction of *isg15* mRNA after injury, lacking the *isg15* upregulation at 72 hpi observed in the wild-type zebrafish (Fig. 2B). As an orthogonal method to confirm that our mutant disrupted interferon signaling in the heart after injury, we visualized the production of *isg15* transcripts using RNAscope at 3 dpi in *ifnphi1^-/-^* and *ifnphi1^+/+^* hearts. This experiment revealed that *ifnphi1^+/+^* zebrafish had enrichment of *isg15* mRNA in the injured zone of their ventricles, while *ifnphi1^-/-^* fish lacked *isg15* enrichment in the injured myocardium (Fig. 2C,D and Supplementary Fig. 1).

**Figure 2:**
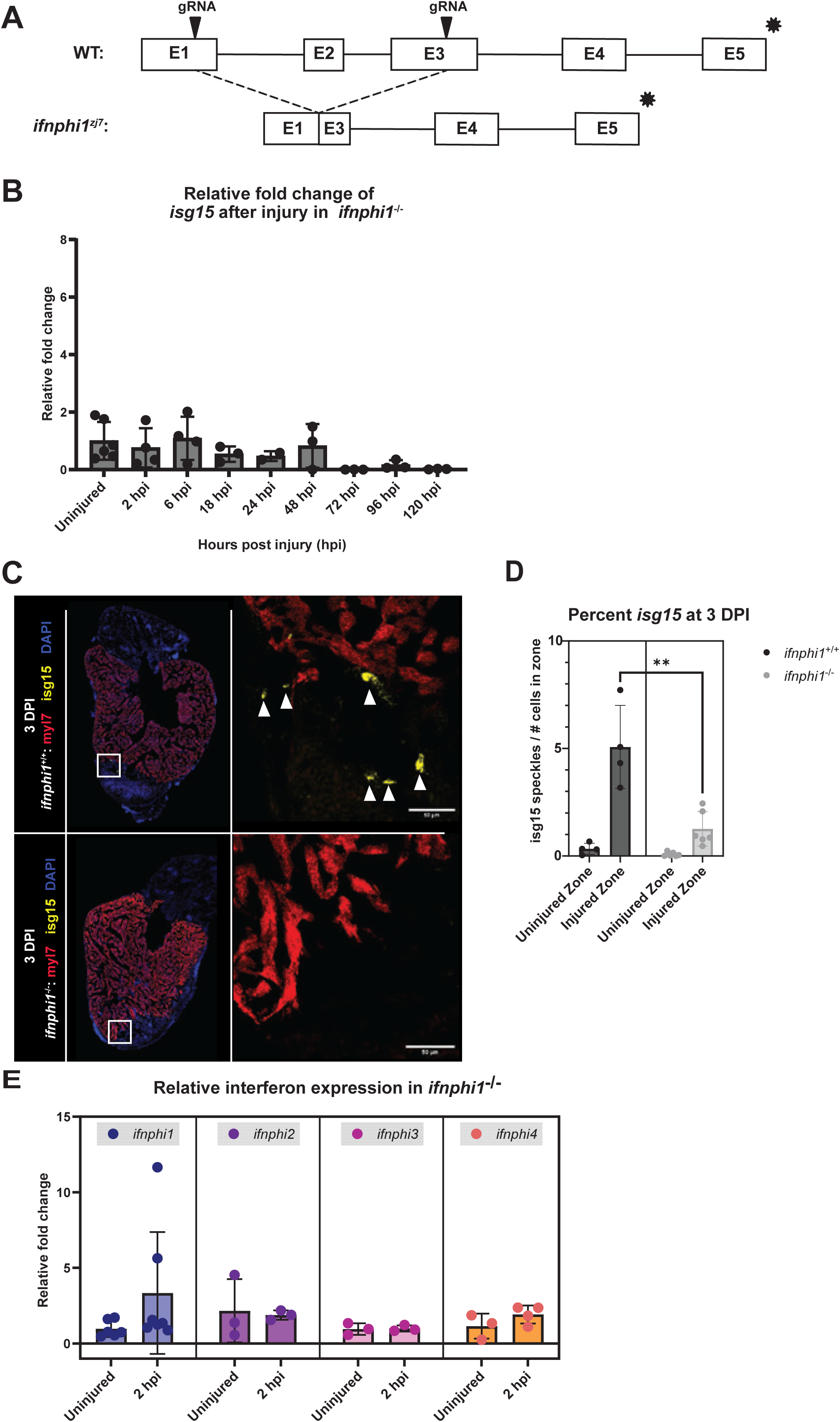
A zebrafish mutant in *ifnphi1* disrupts interferon signaling after heart injury. A) A design for mutating the *ifnphi1* gene, with flanking guide RNAs shown (above) and the *ifnphi*^zj7^ allele with a large deletion that removes part of exon 1, all of exon 2, and part of exon 3 (see Table S1 for sequences). B) A time course of *isg15* gene expression after cryoinjury in *ifnphi1^-/-^* ventricles (2 to 120 hpi) relative to uninjured ventricles. C) Representative images of RNA fluorescence *in situ* hybridization (RNA-FISH) of *isg15* (interferon stimulated gene-15) and *myl7* (labels cardiomyocytes) in wild-type and *ifnphi1^-/-^* ventricle cryosections at 3 dpi. Sections were counterstained with DAPI (blue). D) Percent *isg15* calculated via the number of *isg15* speckles quantified in CellProfiler divided by the number of nucleated cells in the injured and uninjured zones. Welch’s t-test was used to determine statistical significance between the *isg15* speckle percent in the injured zones (p = 0.0023). The dots on the graph represent the uninjured or injured zones of individual ventricles. Data are shown as the mean ±□s.d.; n=4 wild-type and n=6 *ifnphi1^-/-^* ventricles. Scale bar = 50 µm. E) *ifnphi1-4* gene expression in *ifnphi1^-/-^* ventricles at 2 hpi relative to uninjured ventricles.

To ensure that the other Type I interferons were not compensating for the lack of *ifnphi1* in mutants, RT-qPCR was used to test for the upregulation of *ifnphi2-4* in both *ifnphi1^+/+^*and *ifnphi1^-/-^* fish, relative to uninjured *ifnphi1^+/+^*fish (Fig. 2E). Here, *ifnphi2-4* is not upregulated in *ifnphi1^+/+^* or *ifnphi1^-/-^* conditions after injury. Collectively, these experiments provide strong evidence that *ifnphi1* is the primary driver of interferon signaling after cardiac injury in zebrafish, and that *ifnphi1^-/-^* fish have disrupted interferon signaling.

### *ifnphi1* promotes early neutrophil recruitment to the injured myocardium

Given the strong connections between interferon signaling and innate immune cell recruitment, we next examined the role of *ifnphi1* in neutrophil and macrophage recruitment to the injury site. To study the role of interferon signaling in neutrophil recruitment, we labeled neutrophils with the mpx:GFP transgene in wildtype and ifnphi1^-/-^ hearts. We dissected and imaged injured hearts at 1 dpi, when neutrophil recruitment is peaking during heart regeneration (Goumenaki et al., 2024; Julier et al., 2017). Because the impact of cryoinjury can vary between hearts, we calculated the total number of neutrophils and the percentage of neutrophils in the injured and uninjured zones by dividing the total number of neutrophils by the total number of nucleated cells in each injury zone (Fig. 3A,B). By both metrics, *ifnphi1^-/-^* fish had significantly fewer neutrophils in the injury zone than *ifnphi1^+/+^*fish at one day post-injury. To observe if this defect persisted, we conducted the same comparison at 3 dpi. While three days after injury is after the peak neutrophil response in the injury zone, *ifnphi1^+/+^* fish still had a significantly higher number of neutrophils than *ifnphi1^-/-^* fish (Fig. 3C,D). These data suggest that interferon signaling is important for the recruitment of neutrophils during the initial stage of zebrafish heart regeneration.

**Figure 3:**
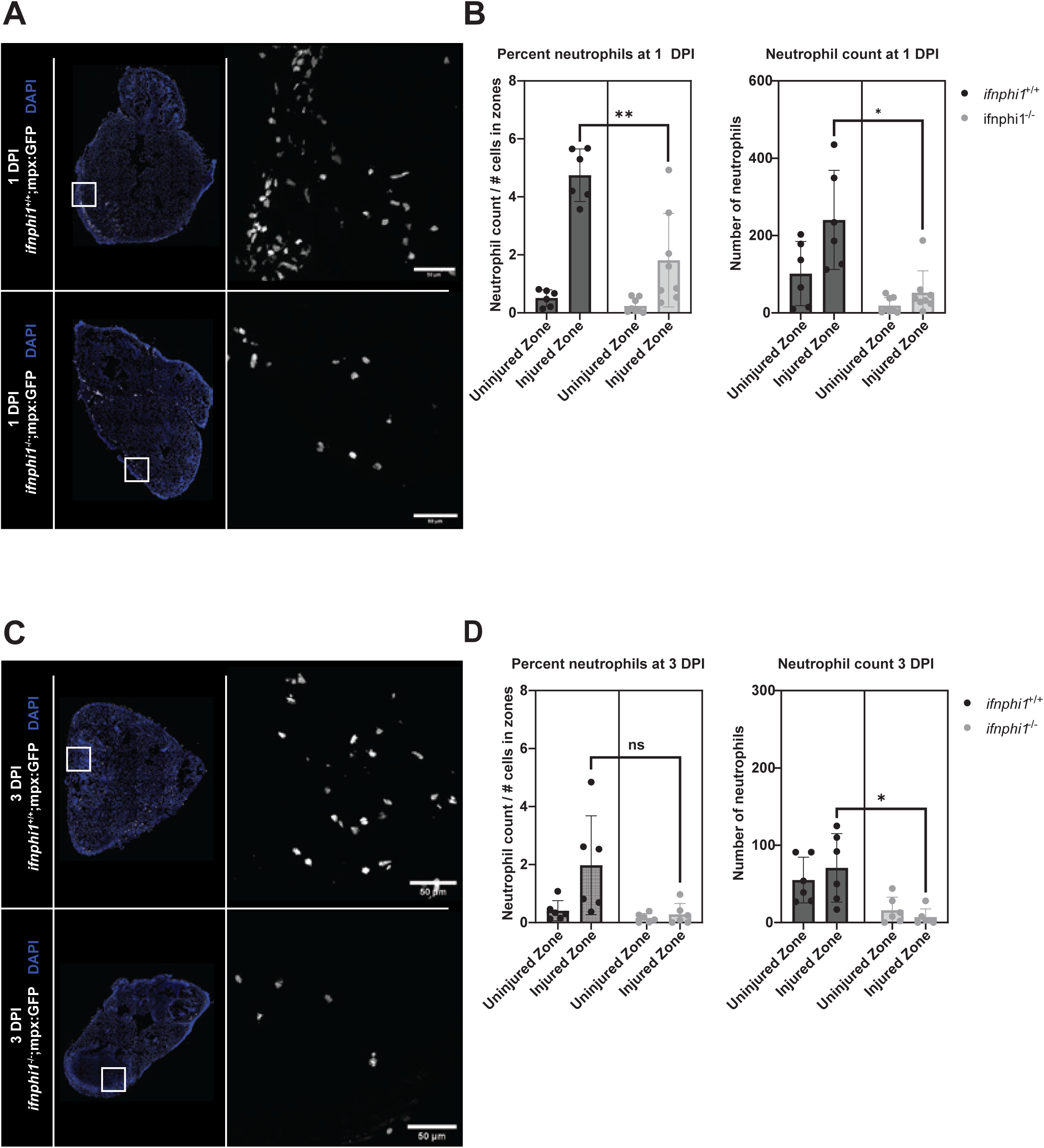
Interferon signaling promotes neutrophil recruitment to injured myocardium. A) Representative images of wild-type Tg(mpx:GFP) and *ifnphi1^-/-^* Tg(mpx:GFP) heart sections at 1 dpi. Sections were counterstained with DAPI (blue). Neutrophils are shown in white. B) Quantification of the percent of neutrophils and absolute number of neutrophils in the injured and uninjured zones of wild-type and *ifnphi1^-/-^* hearts at 1 dpi. The total number of neutrophils was counted in the uninjured and injured zones in wild-type and *ifnphi1^-/-^* hearts (p=0.0137). The percent of neutrophils was calculated by dividing the number of neutrophils by the number of nucleated cells in the uninjured and injured zones in wild-type and *ifnphi1^-/-^* hearts (p=0.0012). C) Representative images of wild-type Tg(mpx:GFP) and *ifnphi1^-/-^* Tg(mpx:GFP) heart sections at 3 dpi. Sections were counterstained with DAPI (blue). D) Quantification of the percent of neutrophils (p=0.0590) and absolute number of neutrophils (p=0.0157) in the injured and uninjured zones of wild-type and *ifnphi1^-/-^* hearts at 1 dpi was calculated as above. Each dot on the graphs represents the uninjured or injured zones calculated from individual sections of distinct heart ventricles. Data are shown as the mean ±□s.d.; n=6 wild-type and n=6 *ifnphi1^-/-^* for one dpi, and n=6 wild-type and n=6 *ifnphi1^-/-^* for 3 dpi. Statistical tests represent Welch’s *t*-test. Scale bar = 50 µm.

### Interferon signaling is dispensable for macrophage recruitment and neovascularization after heart injury

Neutrophils directly interact with macrophages by secreting chemokines and cytokines, thereby influencing the pro-repair, anti-inflammatory identity of infiltrating macrophages (Bouchery & Harris, 2019; Wilgus et al., 2013). In the wound, macrophages phagocytose cellular debris and dying neutrophils, contributing to the transition to an anti-inflammatory immune response (Bouchery & Harris, 2019; Marwick et al., 2018). In addition to shaping the inflammatory response, macrophages closely associate with protruding cardiomyocytes at the border injury zone and help remodel the extracellular matrix to facilitate scar resolution (Constanty et al., 2025). To estimate the number of macrophages that infiltrated the injury site in wildtype and ifnphi1^-/-^ mutant hearts, we measured *mpeg* expression, a classic marker of macrophages, using RNA-FISH. We found that both genotypes have similar amounts of mpeg expression (Fig. 4A,B and Supplementary Fig. 2) suggesting that macrophage abundance was not impacted by the lack of interferon signaling in the mutants. Despite the implications of macrophage recruitment and TLR signaling demonstrated in a previous medaka study, our data suggest that *ifnphi1*-driven interferon signaling does not play a role in macrophage recruitment in zebrafish (Lai et al., 2017) (Fig. 4A,B).

**Figure 4:**
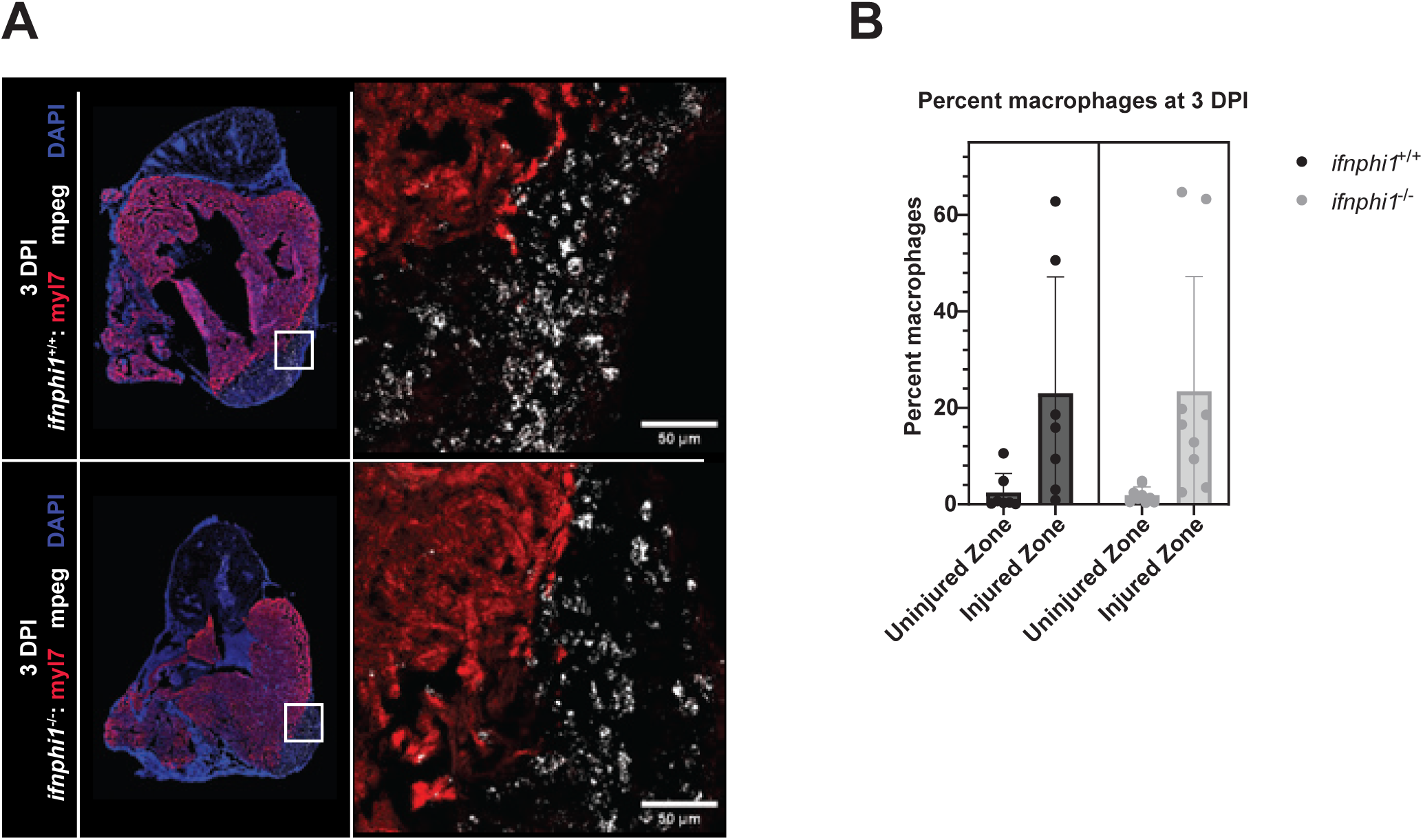
Interferon signaling is not required for macrophage recruitment to the heart injury zone. A) Representative images of RNA-FISH of *mpeg* (labels macrophages) and *myl7* (labels cardiomyocytes), in wild-type and *ifnphi1^-/-^* ventricle cryosections at 3 dpi. Sections were counterstained with DAPI (blue). B) Percent macrophages were calculated by dividing the number of *mpeg* speckles by the number of nucleated cells in the injured and uninjured zones. The dots on the graph represent the uninjured or injured zones calculated from individual sections of distinct heart ventricles. Data are shown as the mean ±□s.d.; n=7 wild-type and n=9 *ifnphi1^-/-^*. Scale bar = 50 µm.

Neovascularization also occurs during the inflammatory response, beginning as early as 15 hpi (Marín-Juez et al., 2016). Previous studies suggest that neutrophils play a role in initiating neovascularization in the wounded myocardium (El-Sammak et al., 2022; Gong & Koh, 2010; Lörchner et al., 2023). We measured neovascularization at 3 dpi, 7 dpi, and in uninjured *ifnphi1^+/+^* and *ifnphi1^-/-^* hearts using RNA-FISH for *kdrl* expression (Fig. 5A-H). The *kdrl* speckles were counted in both injured and uninjured zones of the myocardium and normalized to the number of nucleated cells in each zone (Fig. 5 B,D,G). The enrichment of *kdrl* expression in the injured versus uninjured zones was calculated for each heart (Fig. 5E,H). Additionally, the number of *kdrl* speckles in the uninjured and injured zones for each heart was totaled (Supplementary Fig. 3). We found that uninjured *ifnphi1^+/+^*and *ifnphi1^-/-^* hearts had very similar *kdrl* expression (Fig. 5A,B). Furthermore, at both 3 and 7 dpi, there were no significant differences in *kdrl* expression or enrichment between *ifnphi1^+/+^*and *ifnphi1^-/-^* hearts (Fig. 5C-H). These data suggest that neovascularization in response to heart injury does not require interferon signaling.

**Figure 5:**
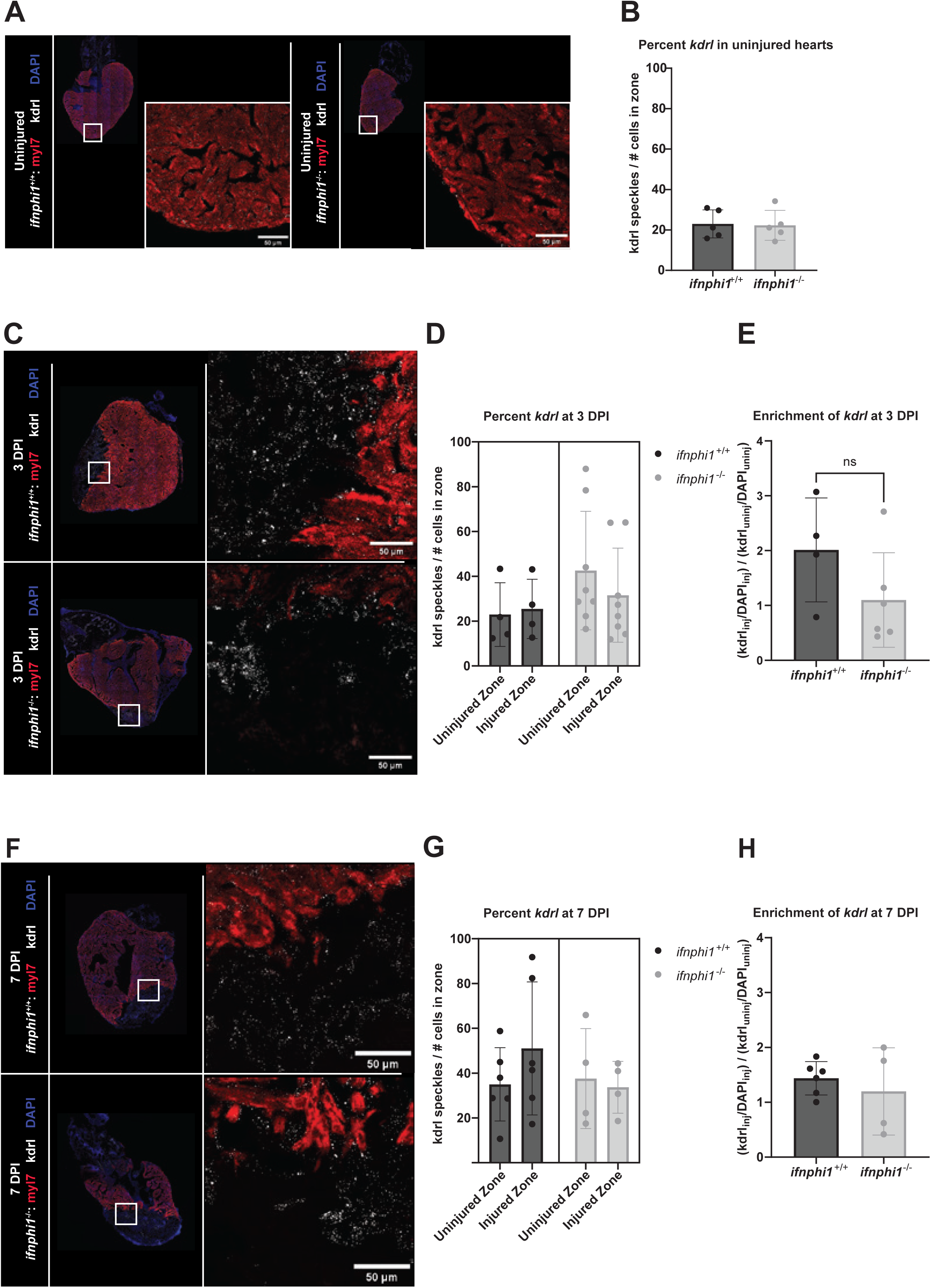
Interferon signaling is not required for neovascularization after heart injury. A, C, F) Representative images of RNA-FISH of *kdrl* (labels vasculature) and *myl7* (labels cardiomyocytes) in uninjured, 3 dpi, and 7 dpi wild-type and *ifnphi1^-/-^* ventricle cryosections. Sections were counterstained with DAPI (blue). B, D, G) Percent *kdrl* was calculated by dividing the number of *kdrl* speckles by the number of nucleated cells in the injured and uninjured zones at each time point. E, H) *kdrl* enrichment was calculated by dividing the normalized *kdrl* speckle count in the injured zone by the normalized *kdrl* speckle count in the uninjured zone of the same heart section, for each heart. The dots on the graph represent the uninjured or injured zones calculated from individual sections of distinct heart ventricles. Data are shown as the mean ±□s.d.; n=5 wild-type and n=5 *ifnphi1^-/-^* for uninjured samples, n=4 wild-type and n=7 *ifnphi1^-/-^* for 3 dpi samples, n=6 wild-type and n=4 *ifnphi1^-/-^* for 7 dpi samples. Scale bar = 50 µm.

### Interferon signaling is not required for scarless regeneration

Immune signaling is critical for scar resolution and scarless regeneration. When innate immune cells are depleted or removed from the inflammatory response to heart injury, collagen persists in the wounded area as a thick, irresolvable scar (Lai et al., 2017; Peterson et al., 2024). We investigated whether interferon signaling is necessary for successful heart regeneration at later time points. After heart cryoinjury, adult zebrafish typically resolve their scar between 30 and 60 dpi. We used Acid Fuchsin Orange-G (AFOG) staining to visualize the extent of remaining scar tissue in *ifnphi1^+/+^* and *ifnphi1^-/-^* hearts at 30 and 60 dpi (Fig. 6A,B and Supplementary Fig. 4 and 5). We identified the most scarred heart section and quantified the size of the remaining scar by normalizing the area of the collagen staining (blue) to the area of the uninjured myocardium (yellow) in each injured heart (Fig. 6C,D). For the 30 dpi samples, the percent collagen in the *ifnphi1^+/+^* and *ifnphi1^-/-^* injured hearts was similar (Fig. 6C), with somewhat more severe injuries in the wild-type hearts. Most hearts had healed from the injury, evidenced by a thicker cortical cardiomyocyte layer and minimal collagen near the injury site. At 60 dpi, the recovery continued in the population of *ifnphi1^+/+^* hearts, but stalled in the population of *ifnphi1^-/-^* hearts.

**Figure 6:**
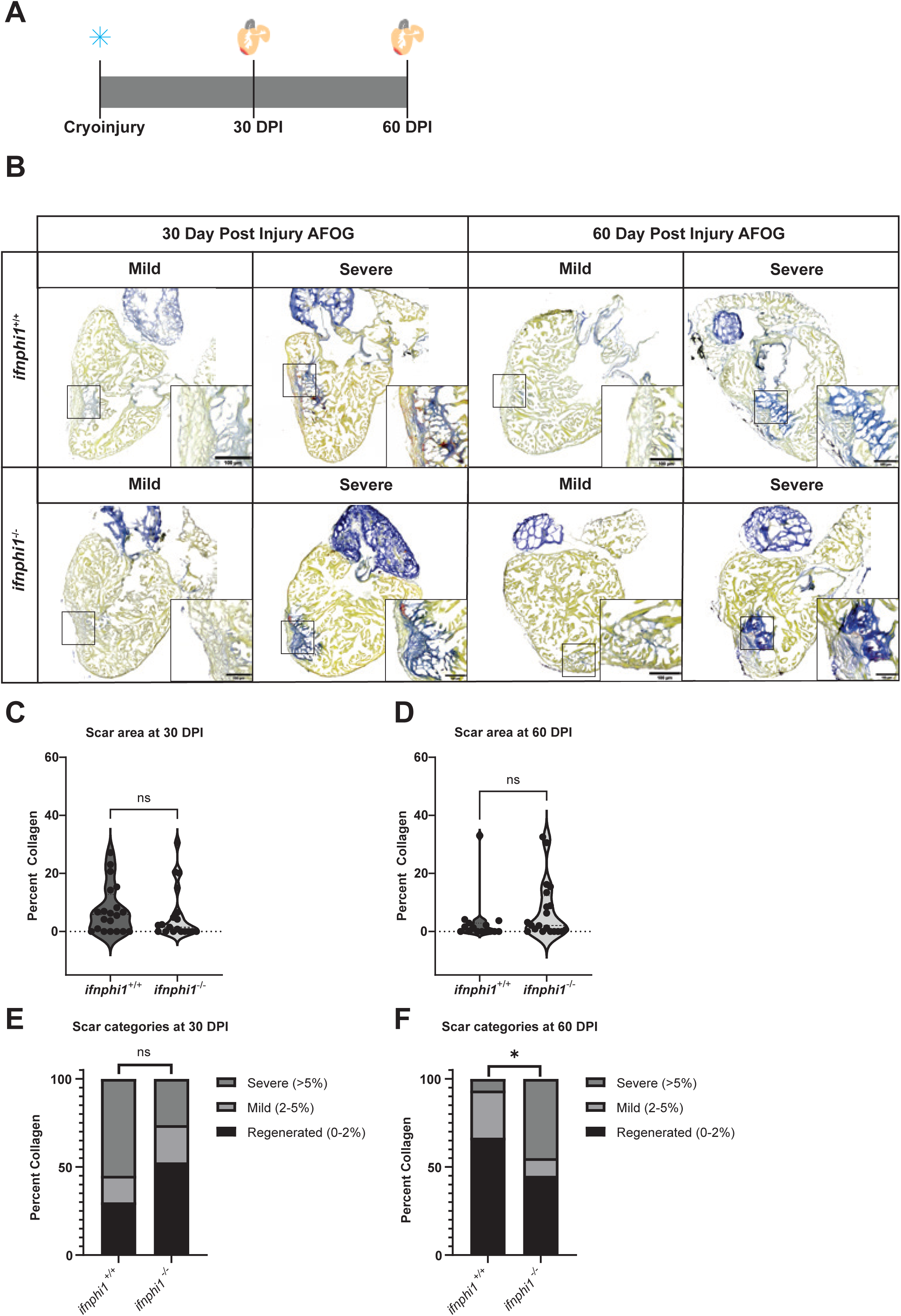
*ifnphi1* mutant hearts are modestly impaired at resolving scars after injury. A) Experimental design for heart injury and collection at 30 and 60 dpi. B) Images of AFOG-stained sections of wild-type and *ifnphi1^-/-^* hearts at 30 and 60 dpi, showing representatives of mild and severe categories of scar resolution. C,D) Quantification of scar size (See Methods) in AFOG-stained hearts between wild-type and *ifnphi1^-/-^* stained sections at each time point. (E,F) Scar severities for all 30 dpi and 60 dpi injured hearts were categorized into three groups: fully regenerated (0-2% collagen), mildly scarred (2-5% collagen), or severely scarred (greater than 5% collagen). Chi-squared test for categorical data showed that 30 dpi categories between wild-type and *ifnphi1^-/-^* are not significantly different (p=0.1517), while 60 dpi categories between wild-type and *ifnphi1^-/-^* are modestly statistically significant (p=0.0378). The dots on the graph represent the uninjured or injured zones calculated from individual sections of distinct heart ventricles. Data are shown as the mean ±□s.d.; n=19 wildtype and n=20 *ifnphi1^-/-^* for 30 dpi samples, n=15 wildtype and n=20 *ifnphi1^-/-^* for 60 dpi samples. Scale bars = 50 µm.

The variation within and between groups at 60 dpi suggested a difference in healing trajectories between the *ifnphi1^+/+^* and *ifnphi1^-/-^* injured hearts. To further investigate these apparent disparities, the injured hearts were categorized based on the percentage of collagen scar as calculated above, representing fully regenerated (0-2% collagen), mildly scarred (2-5% collagen), and severely scarred (>5% collagen) (Fig. 6E,F). This categorical analysis confirmed that at 30 dpi, more *ifnphi1^+/+^* hearts were severely scarred than *ifnphi1^-/-^* hearts, although this difference was not statistically significant (p=0.1517). By 60 dpi, 95% of injured *ifnphi1^+/+^* hearts were either fully regenerated or mildly scarred, as expected (Fig 6E,F). However, regeneration in the *ifnphi1^-/-^* hearts did not follow the same progression in recovery. Nearly half of the mutant hearts remained severely scarred at 60 dpi. The chi-squared statistical test revealed a modestly significant distinction between *ifnphi1^+/+^* and *ifnphi1^-/-^* heart regeneration at this time point (p=0.0378). In summary, our data suggest that interferon signaling has a modest role in resolving the scar during zebrafish heart regeneration.

## DISCUSSION

The inflammatory response to zebrafish heart injury clears cellular debris and promotes scar formation and resolution. Our study characterizes the role of Type I interferon signaling following cardiac injury in zebrafish, revealing its critical and specific contribution to early inflammation. We found that IFN 1, produced by the *ifnphi1* gene, is the primary injury-responsive interferon in the heart, and directs upregulation of ISGs. This response is specific to heart injury, and not sham surgery alone, suggesting that *ifnphi1* is activated by damage-associated molecular patterns (DAMPs) released from dying cells in the injured myocardium rather than pathogen-associated molecular patterns (PAMPs). This aligns with emerging evidence of non-canonical roles for interferon signaling in tissue repair (Cheng et al., 2008; Lai et al., 2017; Leibowitz et al., 2021). The absence of an endogenous interferon response in injured medaka, which cannot regenerate after heart injury, further highlights its potential importance in creating a pro-regenerative environment (Carey et al., 2024; Lai et al., 2017).

We found that interferon signaling is required to attract neutrophils to the injury site early in zebrafish heart regeneration. Roles for neutrophils have been identified in several tissue regeneration contexts. In mouse skin and spinal cord injuries, when neutrophils were depleted, adult mice exhibited slower wound healing (Nishio et al., 2008; Stirling et al., 2009). Zebrafish neutrophils control epicardial cell expansion just days after heart injury (Peterson et al., 2024). When zebrafish neutrophils were depleted, there was significantly less epicardial cell expansion and less cardiomyocyte proliferation; however, the long-term effects of neutrophil depletion were not studied. Here, we found a correlation between interferon signaling, neutrophil recruitment, and timely scar resolution during zebrafish heart regeneration.

We also found that disrupting interferon signaling did not impact macrophage recruitment to the site of injury or interfere with the re-growth of new blood vessels. We suspect that other pro-inflammatory cytokines, independent of the interferon signaling pathway, were likely sufficient to recruit macrophages and trigger vascular growth in the injured myocardium (Luz et al., 2025; Nguyen-Chi et al., 2017). Interestingly, the artificial induction of interferon signaling in the medaka heart, via Toll-like receptor activation, enhanced macrophage recruitment, not neutrophil recruitment (Lai et al., 2017). This highlights the differences in immune response to injury between species.

We found a mild regeneration disparity between the *ifnphi1^+/+^*and *ifnphi1^-/-^* injured hearts, where regeneration stalls in the absence of interferon signaling. We hypothesize that the early impact on neutrophil recruitment in *ifnphi1^-/-^* hearts may have resulted in slower wound healing. Elucidating the specific mechanisms that connect neutrophil activity with scar resolution remains an exciting avenue for study. Future work could interrogate these mechanisms using mutants or small-molecule inhibitors of neutrophil activity (Isles et al., 2019; Kell et al., 2018; Peterson et al., 2024). Despite reduced neutrophil infiltration in all mutant hearts, approximately half of *ifnphi1^-/-^* fish were still able to regenerate completely by 60 dpi. This suggests that the inflammatory process, including phagocytosis of cellular debris and remodeling of the extracellular matrix to facilitate regenerative scarring, was adequately executed in the absence of interferon signaling in these individuals. This variability between individuals may have many underlying reasons, including variation in the intensity of injury or more subtle differences in the tempo of damage sensing, cell recruitment, or tissue regrowth.

Collectively, this study underscores the concept of regenerative robustness, where redundant mechanisms protect outcomes to the function of the organ and organism. Understanding how these mechanisms drive zebrafish heart regeneration may reveal novel therapeutic targets to promote repair in non-regenerative mammalian hearts, where key compensatory pathways may be lost or insufficient.

## MATERIAL AND METHODS

### Fish husbandry and zebrafish lines

All zebrafish lines were maintained in accordance with standard husbandry practices at the CBRZ zebrafish facility at the University of Utah This study was conducted with approval of the Office of Institutional Animal Care and Use Committee (IACUC no. 20-07013) of the University of Utah’s animal care and use program.

### Generation of a zebrafish *ifnphi1* mutant line

To generate a stable *ifnphi1* mutant line, *ifnphi1* was targeted using CRISPR-Cas9 gene editing. 4 guide RNAs (gRNAs) targeting exons 1-3 of the *ifnphi1* gene were designed and ordered from IDT. To generate the crRNA/gRNA duplex for each gRNA, a 100uM solution of crRNA and tracrRNA was made for each duplex in IDT gRNA buffer. Equal volume of tracrRNA (5 μl) and gRNA (5 μl) was combined annealed via PCR to generate the crRNA-gRNA duplex for each gRNA. The PCR program to anneal crRNA and gRNA is as follows: 95°C, 5 min; cool at 0.1°C/s to 25°C; 25°C, 5 min; cool to 4°C rapidly.

Along with these gRNAs (800 ng/ µl), SpCas9 protein (from IDT), and phenol red were coinjected into WT embryos. Total volumes: crRNA/gRNA 1-4 (1.24 µl each), SpCas9 (0.5 µl at 10 µg/µl), and phenol red (1 µl). 1 nanoliter was injected into each embryo. Mosaic embryos were raised to adulthood and incrossed with each other before being outcrossed to wild-type fish. Primers amplifying the CRISPR-editing region were used to select outcrossed fish with large deletions. Specifically, embryos from this outcross carry large deletions, ∼800 bp. To establish a homozygous mutant line, these heterozygous mutant fish were incrossed. Three primers were used to distinguish between homozygous mutants, heterozygous fish, and wild-type fish in two separate PCR reactions (Supplementary Table 1). All experiments presented compare homozygous mutant zebrafish with WT siblings. This line was given the designation *ifnphi^zj7^*. Mutant zebrafish were crossed into the Tg(mpx:GFP) line to label neutrophils (Buchan et al., 2019).

### Cardiac cryoinjury

Cryoinjuries were performed on the ventricular apex of anesthetized zebrafish as described previously (González-Rosa et al., 2011). 0.02% Tricaine (MS-222) was used to anesthetize zebrafish. Zebrafish were positioned on a moist sponge. A small thoracic incision was made with forceps and dissecting scissors to expose the ventricular apex. Using a liquid nitrogen-chilled copper wire cryoprobe (0.5 mm in diameter, as previously described (González-Rosa & Mercader, 2012)), the ventricular apex was injured with the probe for 23 seconds. Following injury, the fish were revived in freshwater tanks and returned to the facility for monitoring.

### Histological methods

Hearts were fixed in 4% paraformaldehyde (PFA, Electron Microscopy Sciences) in 1X PBS for 24 hours, then sucrose-treated sequentially from 10% to 30% sucrose. Sucrose-treated hearts were then mounted in Optimal Cutting Temperature (O.C.T., Fisher HealthCare) embedding medium and frozen at -80 C. Hearts from Tg(mpx:GFP) animals were sectioned at 12 µm and mounted in Fluoroshield with DAPI (Sigma-Aldrich). For Acid Fuchsin Orange G (AFOG, Sigma) staining, O.C.T. mounted hearts were sectioned at 7 µm. Bouin’s solution (Sigma Aldrich) was used to fix heart tissue before staining with PT/PM (phosphotungstic acid/phosphomolybdic acid, Sigma-Aldrich) and AFOG stain. After staining, sections were washed twice with 0.5% acetic acid (30 seconds per wash), four times with 100% ethanol (1 minute per wash), and three times with xylenes (5 minutes per wash). For AFOG imaging, the ECHO Revolution 2.0 confocal microscope was used to capture brightfield images at 10X.

### RNAscope *in situ* hybridization

Zebrafish hearts were fixed for 24 hours in 4% paraformaldehyde (PFA) in 1X PBS, sequentially sucrose-treated (10% to 30%), and O.C.T. embedded. Hearts were then cryosectioned at 12 µm. RNAscope (Advanced Cell Diagnostics, Hayward, CA, USA) was performed using the RNAscope® Multiplex Fluorescent Detection Kit v2 protocol, with previously published probes (Carey et al., 2024). The Zeiss 880 confocal microscope was used to capture RNAscope images at 40X.

### Imaging quantification

ImageJ was used to analyze images of heart sections probed for *myl7* (cardiomyocytes), *mpeg1.1* (macrophages), *isg15* (interferon-stimulated gene-15), *kdrl* (vascular), and DAPI. To separate the wound from the uninjured ventricle, we set a threshold on the cardiomyocyte channel to create an uninjured ventricle mask. Using the DAPI channel and the difference between the uninjured ventricle mask and the DAPI mask, an injured area mask was created for every heart. Masked images of each probe in the injured and uninjured areas were uploaded to CellProfiler software (version 4.2.1) using a modified CellProfiler ‘Speckle Counting’ pipeline (Stirling et al., 2021). Total number of nucleated cells in the DAPI channel and *mpeg1.1, isg15*, and *kdrl* speckles were identified using the IdentifyPrimaryObjects module. Macrophages and nuclei were related using the RelateObjects CellProfiler module. The percent of macrophages relative to the total number of cells in injured and uninjured areas was calculated. Both *isg15* and *kdrl* speckles were counted regardless of the nuclei relationship. Percent *kdrl* and *isg15* speckles relative to the total number of cells in injured and uninjured areas was calculated. Furthermore, the enrichment of *kdrl* expression in the injured versus uninjured section was calculated for each heart.

For Tg(mpx:GFP) neutrophil counts in injured wildtype and interferon 1Δ fish at 1- and 3-dpi, uninjured and injured masks were created using ImageJ as described above. A similar CellProfiler ‘Speckle Counting’ pipeline was used to count the number of neutrophils in the injured and uninjured areas.

For scar quantification in AFOG images, ImageJ software was used. Here, the scar size was calculated by measuring the collagen (blue) stained area and divided by the total myocardium, and multiplied by 100 to generate a percentage.

### RT-qPCR

RNA was purified from injured ventricles using PicoPure (Thermo Fisher) and converted to complementary DNA (cDNA) using QuantiTect Reverse Transcription Kit, using the included RT primer mix (Qiagen). cDNA was amplified using BrilliantII SYBR Green QPCR Master Mix (Agilent) with 10 pmol of each primer. Quantitative reverse-transcription PCR (qRT-PCR) was performed using the QuantStudio 5 (Thermo Fisher Scientific) protocol. Elongation factor 1-alpha (*ef1*α) expression was measured for each biological triplicate in every experiment. The ΔCt method was used to analyze the data, with *ef1*α *as the* normalization factor for each sample. The fold change was calculated using the ΔΔCt method, where 2-ΔΔCt of injured samples is relative to the average of 2-ΔΔCt of uninjured heart samples.

### Statistical analysis

Statistical analyses for column and nested data were performed using GraphPad Prism (v.10.5.0). Data distribution was assessed using the Shapiro-Wilk normality test. The significance level was set to p=0.05, and the standard deviation from the mean is indicated on the figures. To determine the statistical significance of the normal data, Welch’s t-test was used.

For the categorical scar analysis (Figure 6E,F), a chi-squared test statistic was calculated for both the 30 dpi and 60 dpi datasets. Then, using this chi-squared test statistic and degrees of freedom=2, a p-value was derived.

---

**Supplementary Figure 1: *isg15* speckle count in *ifnphi1^-/-^* versus wild-type injured ventricles at three dpi.** The number of *isg15* speckles counted using the CellProfiler pipeline is represented in the injured and uninjured zones of each fish at 3 dpi. Each dot represents the injured and uninjured zones of one ventricle. Differences in *isg15* expression in the injured zones between *ifnphi1^-/-^* and *ifnphi1^+/+^*were significant with *p*=0.0141 using Welch’s t-test.

**Supplementary Figure 2: *mpeg* speckle count in *ifnphi1^-/-^* versus wild-type injured ventricles at three dpi.** The number of *mpeg* speckles counted using the CellProfiler pipeline is represented in the injured and uninjured zones of each fish at 3 dpi. Each dot represents the injured and uninjured zones of one ventricle. Differences in *mpeg* expression in the injured zones between *ifnphi1^-/-^* and *ifnphi1^+/+^* were not significant with *p*=0.2119 using Welch’s t-test.

**Supplementary Figure 3: *kdrl* speckle count in *ifnphi1^-/-^* versus wild-type injured ventricles.** A-C) The number of *kdrl* speckles counted using the CellProfiler pipeline is represented in the injured and uninjured zones of each fish at 3 and 7 dpi. Each dot represents the injured and uninjured zones of one ventricle. Differences in *kdrl* expression in the injured zones between *ifnphi1^-/-^* and *ifnphi1^+/+^*were not significant at any post-injury time point.

**Supplementary Figure 4: All AFOG images at 30 dpi.** A) All *ifnphi1^+/+^*hearts 30 dpi AFOG stained. B) All *ifnphi1^-/-^* hearts 30 dpi AFOG stained. Scale bars = 500 µm.

**Supplementary Figure 5: All AFOG images at 60 dpi.** A) All *ifnphi1^+/+^*hearts 60 dpi AFOG stained. B) All *ifnphi1^-/-^* hearts 60 dpi AFOG stained. Scale bars = 500 µm.

**Supplementary Table 1:** Sequence information for the *ifnphi1^zj7^*allele, *ifnphi1^zj7^* genotyping primers, RT-qPCR primer sequences, and gRNA sequences.

## ACKNOWLEDGMENTS

We thank all members of the Gagnon laboratory for their assistance in completing this project, especially Drew Janik, Jenna Weber, and Kate Scuderi, for their contributions to surgical procedures, Connor Shrader for advice on statistics, and Hailey Hollins and Clay Carey for advice and technical support. We acknowledge the staff of the Centralized Zebrafish Animal Resource facility, ARUP Research Histology Core Laboratory for sectioning assistance, and the Cell Imaging Core at the University of Utah, specifically Xiang Wang. We thank Matthew Mulvey for the use of his microscope. These facilities did not provide financial support.

## FUNDING

This project was supported by National Institutes of Health grants T32HD007491 (A.V.S.) and R35GM142950 (J.A.G.), and by startup funds from the University of Utah Henry Eyring Center for Cell and Genome Science (J.A.G.).

## CONTRIBUTIONS

Conceptualization, methodology, software, formal analysis, and investigation: A.V.S.; Writing of the original draft: A.V.S.; Writing review and editing: A.V.S. and J.A.G.; Supervision: J.A.G.; Project administration: J.A.G.; Funding acquisition: J.A.G.

## Notes

### Competing Interest Statement

The authors have declared no competing interest.

