## Supplementary figures and images for "Interferon signaling promotes early neutrophil recruitment after zebrafish heart injury"

### Supplemental Figures

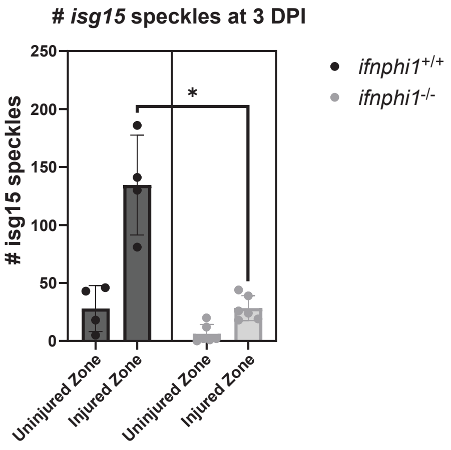

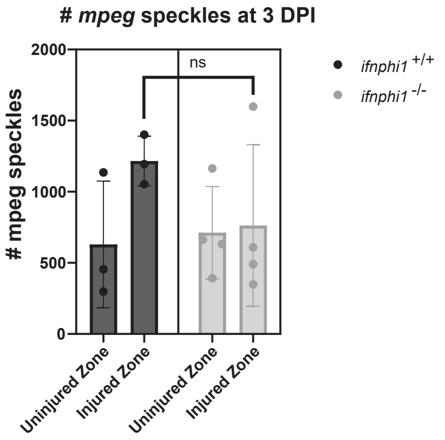

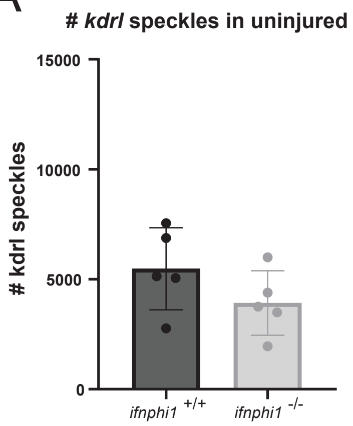

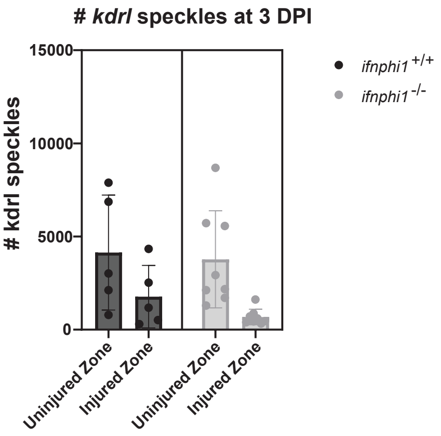

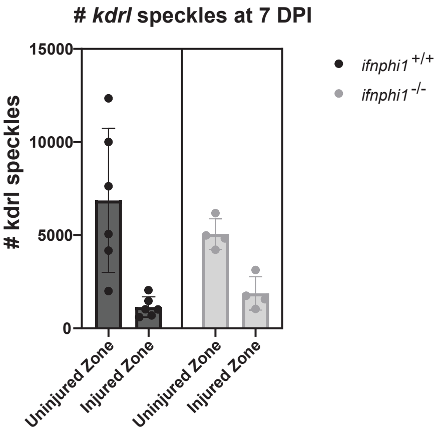

A WT 30 DPI AFOG:

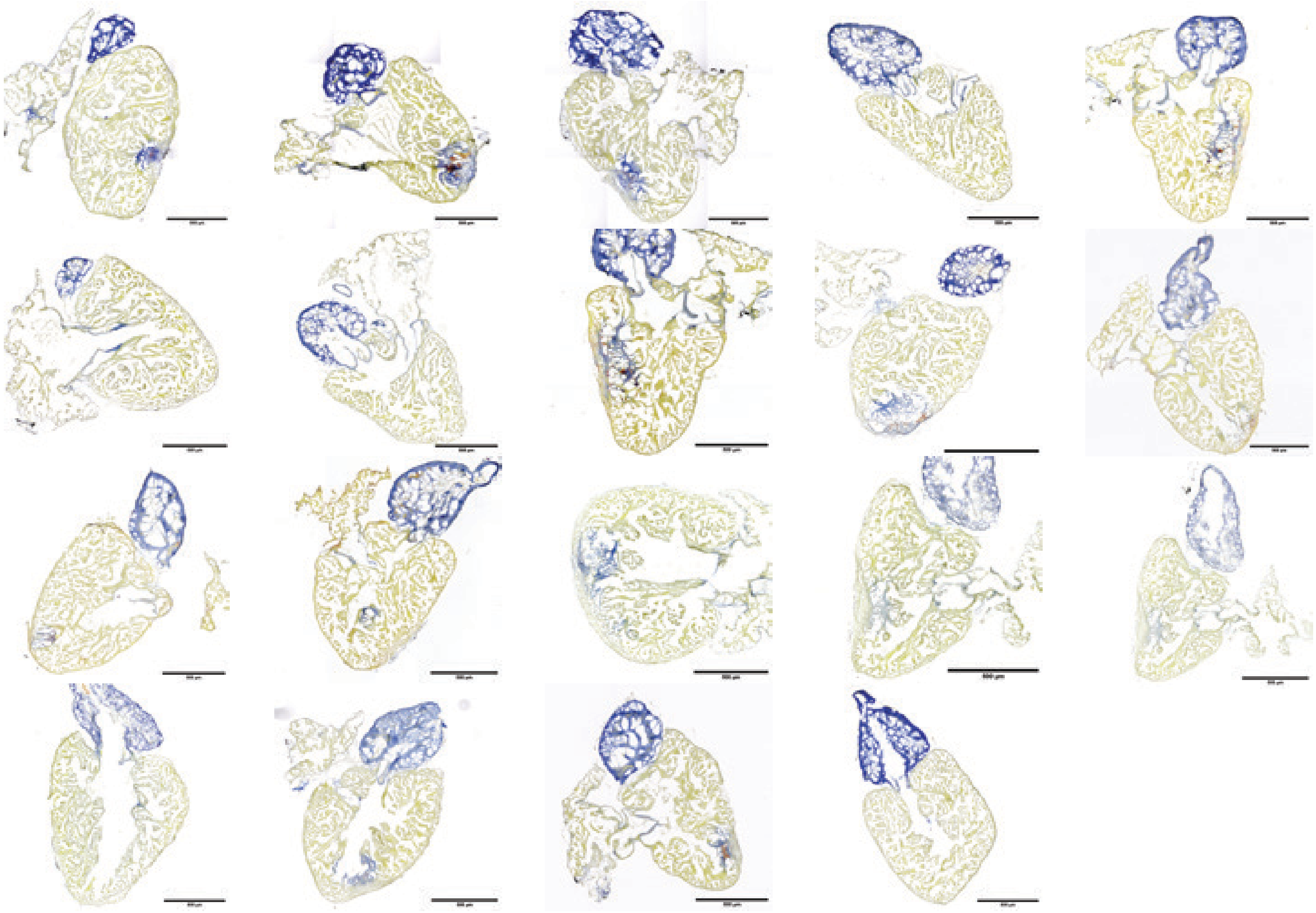

B *ifnphi1*$^{-/-}$ 30 DPI AFOG:

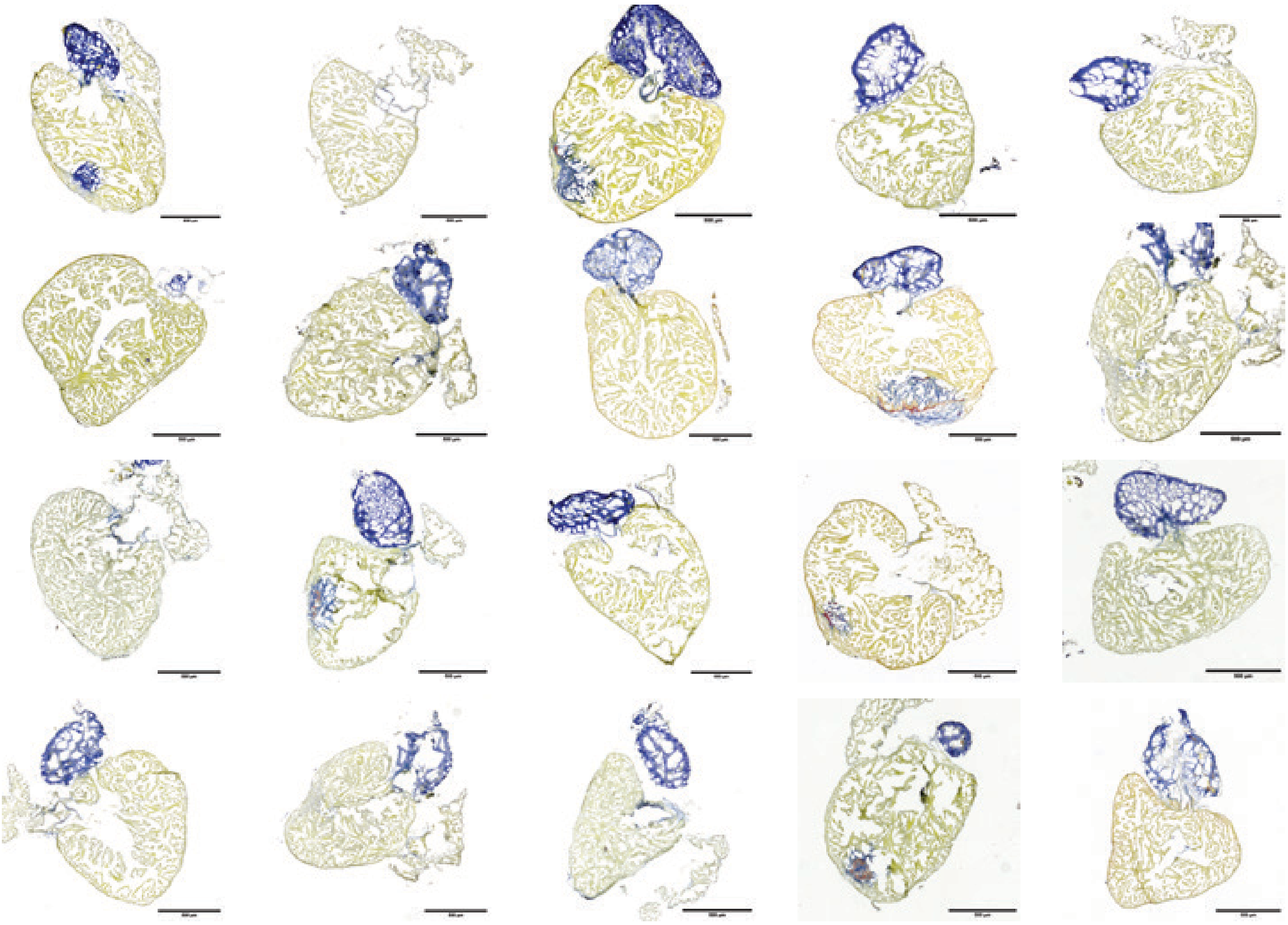

A WT 60 DPI AFOG:

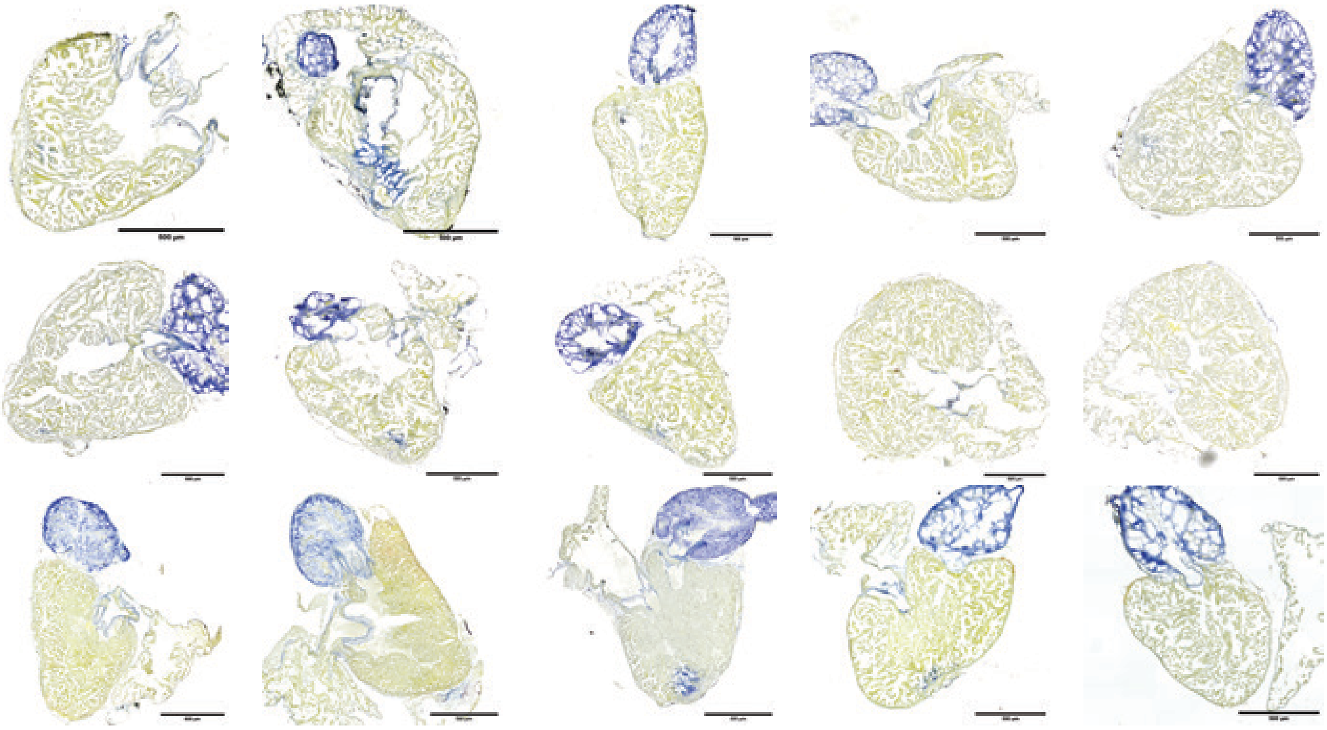

B *ifnphi1*$^{-/-}$ 60 DPI AFOG:

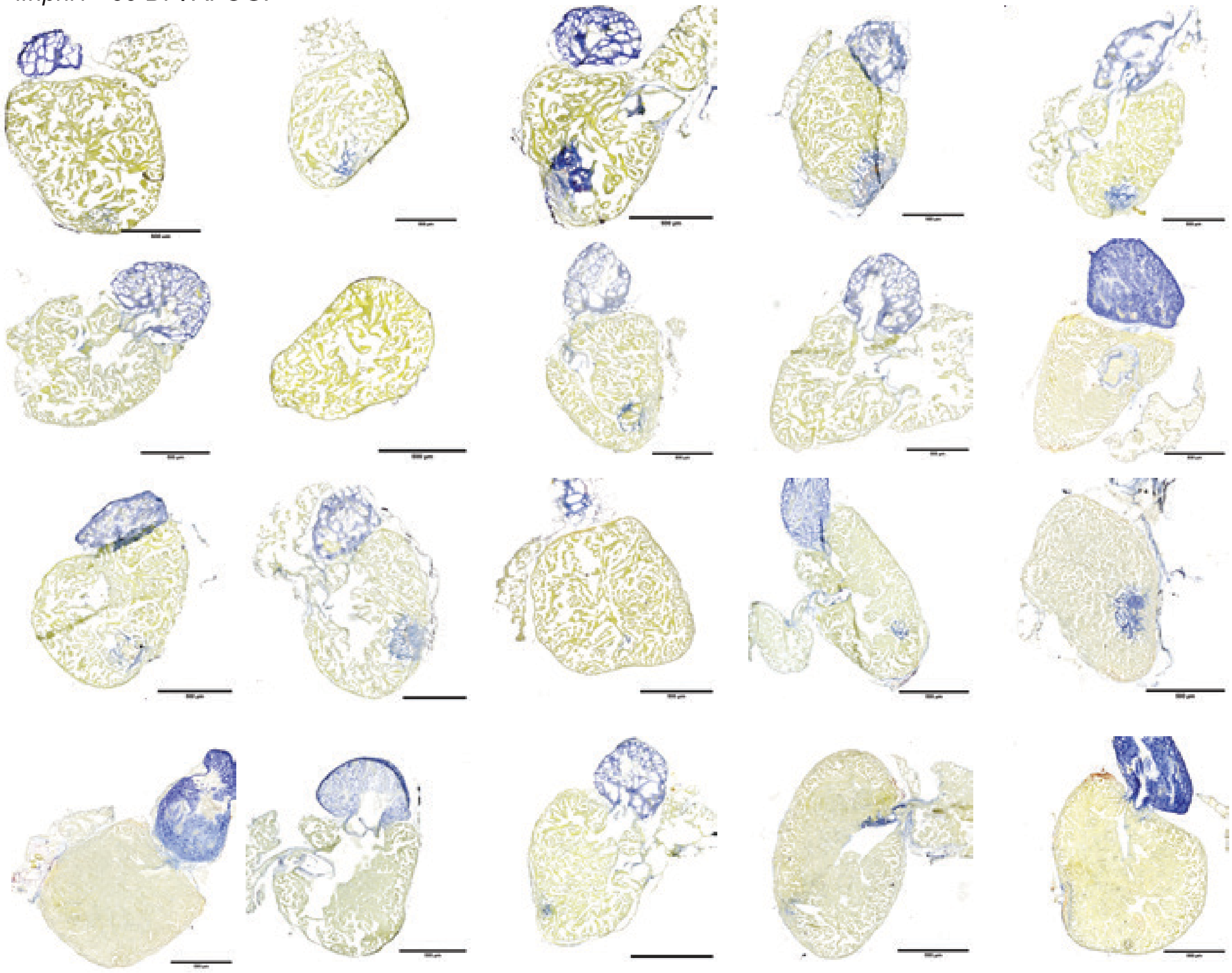
